# Objective Assessment of Beat Quality in Transcranial Doppler Measurement of Blood Flow Velocity in Cerebral Arteries

**DOI:** 10.1101/554568

**Authors:** Kian Jalaleddini, Nicolas Canac, Samuel G. Thorpe, Benjamin Delay, Amber Y. Dorn, Fabien Scalzo, Corey M. Thibeault, Seth Wilk, Robert B. Hamilton

**Author notes:** These authors contributed equally to the work (Kian Jalaleddini, Nicolas Canac).

## Abstract

**Objective:** Transcranial Doppler (TCD) ultrasonography measures pulsatile cerebral blood flow velocity in the arteries and veins of the head and neck. Similar to other real-time measurement modalities, especially in healthcare, the identification of high quality signals is essential for clinical interpretation. Our goal is to identify poor quality beats and remove them prior to further analysis of the TCD signal.

**Methods:** We selected objective features for this purpose including Euclidean distance between individual and average beat waveform, cross-correlation between individual and average beat waveform, ratio of the high frequency power to the total beat power, beat length, and variance of the diastolic portion of the beat waveform. We developed an iterative outlier detection algorithm to identify and remove the beats that are different from others in a recording. Finally, we tested the algorithm on a dataset consisting of more than 16 hours of TCD data recorded from 48 stroke and 35 in-hospital control subjects.

**Results:** We assessed the performance of the algorithm in estimating clinically important TCD parameters by comparison to those identified from beats hand-annotated by an expert. The results show that there is strong correlation between the two that delineates the algorithm has successfully recovered clinically important features. We obtained significant improvement in estimating the TCD parameters using the algorithm accepted beats compared to using all beats (r>0.78, p<0.01).

**Significance:** Our algorithm provides a valuable tool to the clinicians for automated detection of the reliable portion of the data. Moreover, it can be used as a pre-processing tool to improve the data quality for machine learning algorithms for automated diagnosis of pathologic beat waveform.

## I. INTRODUCTION

*Transcranial Doppler* (TCD) ultrasonography is a diagnostic technique for rapid, non-invasive assessment of cerebrovascular health [1]. TCD was first used in neurology in the 1980s and measures *cerebral blood flow velocity* (CBFV) [2]. TCD has demonstrated utility in the diagnosis of clinical conditions such as acute ischemic stroke [3], intracranial pressure [4], [5], sickle cell disease [6], traumatic brain injury [7], dementia [8], cerebral emboli [9] and many others [10], [11].

CBFV beat waveform morphology has not been extensively studied mainly due to the noise level and lack of signal processing techniques developed for TCD signal analysis [4], [12]. As seen in Fig. 1, beat waveform morphology can exhibit a high degree of variability due to noise within the signal often caused by the movement of the subject or sonographer, or the electronics. However, clinically important parameters such as *mean Cerebral Blood Flow Velocity* (mCBFV) or *Pulsatility Index* (PI), which may be used to aid in diagnostic assessment, are often extracted from this same highly variable TCD beat waveform morphology [4], [7], [13], [14]. Thus, being able to reliably identify and exclude the low-quality portions of the signal in a consistent, quantitative manner is imperative for the accurate extraction of clinically relevant TCD parameters and any ensuing clinical interpretation based on those parameters.

**Fig. 1.**
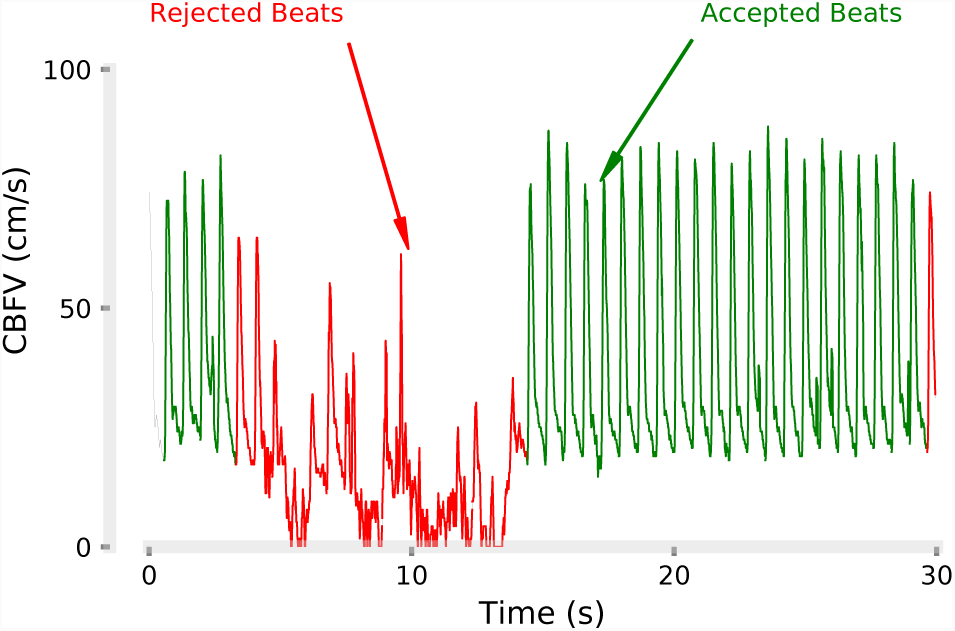
Measurement of *Cerebral Blood Flow Velocity* (CBFV) can exhibit noise within a TCD recording. Currently, an expert may be needed to manually inspect and label the poor-quality beats to allow for removal in subsequent analyses of the beat waveform morphology.

The problem of variability and noise in the CBFV recording is often mitigated by manual classification of beats that “appear” different than others in a recording by an expert. This process can be time-consuming, labor intensive, subjective, and difficult to reproduce. In addition, it precludes real-time computation of clinically relevant parameters. Consequently, there is a need for objective beat quality criteria and an automated algorithm for classification of poor-quality beats to make the analysis comprehensive, reliable, and repeatable. The purpose of this paper is to present candidate features for objective quantification of beat quality, develop a classification algorithm to identify poor-quality beats in a recording, and to validate the resulting tool through comparison to a human expert using a relatively large dataset consisting of healthy and pathological beat waveforms.

## II. METHODS

In this section, we describe the experimental protocol for data acquisition, the features we used to characterize beat quality, and an algorithm to classify and remove low-quality beats.

### A. Experiments

#### 1) Subjects

We acquired TCD waveforms from 83 subjects enrolled in-hospital at Erlanger Health Systems Southeast Regional Stroke Center in Chattanooga, TN. The subjects were either diagnosed with *Large Vessel Occlusion* (LVO) of intracranial arteries confirmed with *Computed Tomography Angiography* (CTA) or in-hospital control subjects who arrived at the hospital presenting with stroke symptoms, but were later confirmed negative for LVO by CTA imaging. We also performed a follow-up TCD recording on the LVO subjects within 72 hours of injury. In total, we acquired data from 149 sessions. The experiment protocols were approved by University of Tennessee College of Medicine Institutional Review Board (ID: 16-097).

#### 2) Recording

A trained technician acquired TCD scans using a 2 MHz hand-held probe. CBFV signals associated with the left or right *Middle Cerebral Arteries* (MCA) were identified by insonating through the transtemporal windows and recording the CBFV at the sampling rate of 125 Hz. The technician obtained recordings for as many depths as possible between 45-60 mm. Once a CBFV signal with a smooth fitting envelope trace was identified and optimized at a specific depth, a recording interval would begin, continuing for a total duration of 30 s. In total, an expert manually inspected 16 hours, 6 minutes, 3 seconds of data that had 76,676 beats of which 8,097 beats (≈ 10%) were rejected.

### B. Theory

#### 1) Features

For each recorded 30 seconds TCD measurement, we identified beat start and end times using an algorithm developed [15], [16] as shown in Fig. 2(A). Then, we normalized individual beat lengths to the median beat length in each measurement. This was achieved by padding the beats whose lengths were shorter than the median beat length with the last values of the CBFV and truncating the beats whose length was longer than the median beat length. Then, from the *l* equi-length normalized beat waveforms *CBFV*_*b1*_(*k*), …, *CBFV*_*bl*_(*k*), we computed the average CBFV beat waveform (see Fig. 2(B)) denoted by 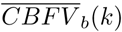 and extracted the following features for each individual beat.

**Fig. 2.**
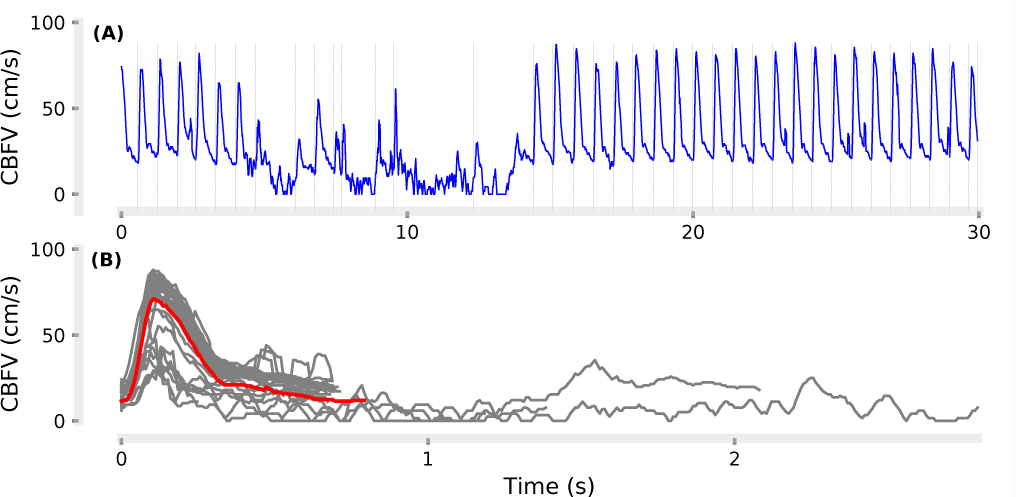
Beat start/end times were identified from measurement of CBFV using our beat detection software (A), and the average beat waveform 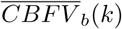 (red beat) was computed from ensemble averaging of individual beat waveforms (grey beats) (B).

##### Euclidean Distance (ED)

*ED*_*i*_ is the Euclidean distance of the *i*-th equi-length beat waveform from 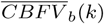:

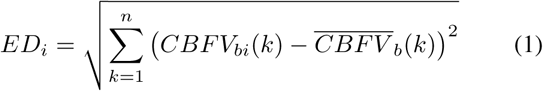

where *n* is the number of samples in the equi-length beats and *i* ∈ {1*, …, l*}

##### Cross-Correlation (CC)

*CC*_*i*_ is the maximum of cross-correlation coefficient between *CBFV*_*bi*_(*k*) of the *i*-th beat and 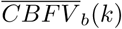:

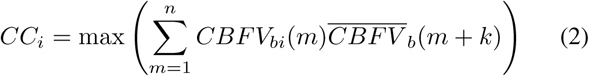

##### Beat Length (BL)

*BL*_*i*_ is the length of the *i*-th original (vs equi-length) beat.

##### High Frequency Noise Power (HFNP)

*HFNP*_*i*_ is the ratio of the high-frequency to low-frequency power for the *i*-th beat.

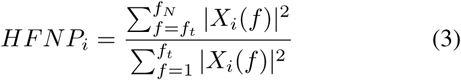

where *X*(*f*) is the discrete Fourier transform of the CBFV, and *f*_*t*_ represents the threshold frequency and *f*_*N*_ is the Nyquist frequency.

##### Diastolic Variance (DV)

*DV*_*i*_ is the variance of the *i*-th CBFV in the diastolic portion of the beat which was defined as the last 20% of the beat.

See Fig. 3 for an illustration of the distributions of these features for a sample recording.

**Fig. 3.**
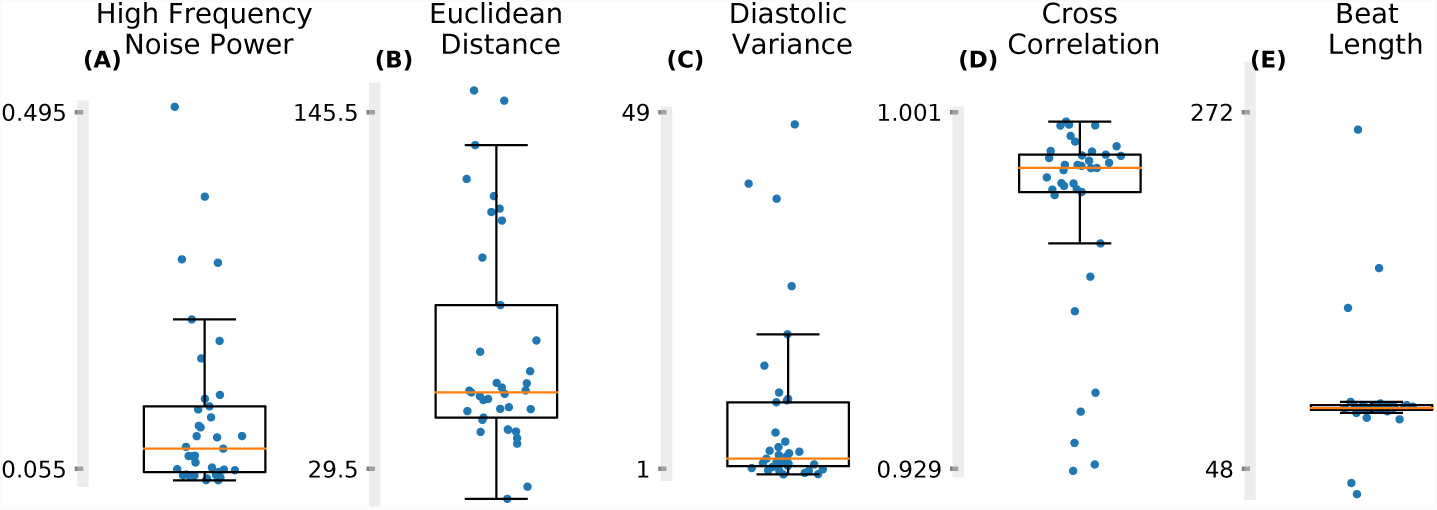
Five features are assembled from individual beat CBFV waveforms: (A) *High-Frequency Noise Power* (HFNP) is the ratio of the high-frequency to low-frequency power; (B) *Euclidean Distance* (ED) is the euclidean distance from the median beat waveform; (C) *Diastolic Variance* (DV) is the variance of the diastolic portion of the beat waveform; (D) *Cross-Correlation* (CC) is the cross-correlation with the median beat waveform; (E) *Beat-Length* (BL) is the beat length in samples.

#### 2) Iterative Outlier Detection Algorithms

We adapted an iterative version of the *InterQuartile Range* (IQR) outlier detection method. The IQR method, first proposed by Tukey [17], [18], works by calculating *IQR* = *q*_3_*-q*_1_ where *q*_1_ and *q*_3_ are the first and third quadrants. It labels as outliers the data points that fall below *B*_*L*_ = *q*_1_ *-*1.5*IQR* or above *B*_*H*_ = *q*_3_ + 1.5*IQR*. The range is shown by the location of the whiskers in the box plot, Fig. 3. The *Iterative IQR* (IIQR) method works by identifying the farthest outlier from the boundary, if any, and removing one outlier at each iteration, continuing until no further outliers are present. This ensures that the boundaries *B*_*L*_ and *B*_*H*_ are not biased because of the presence of large outliers.

*Iterative InterQuartile Range* (IIQR) Algorithm: The following iterative algorithm works by identifying outliers from the set *F* = {*f*_1_, *f*_2_*,, f*_*l*_} for a beat feature described above (e.g. ED) where *n* is the total number of beats.

1. Denote the list of outlier indices as *I*^*o*^ = {}.
2. Calculate *q*_1_ and *q*_3_ that are the 25-th and 75-th percentiles of *F.*
3. Populate *F* ^*o*^ with the outliers:

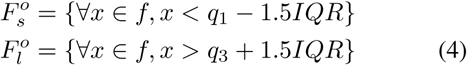 If 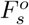 and 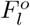 are empty, then exit, returning the list of the outlier indices *I*^*o*^.
4. Find the outlier that is the farthest from the boundaries:

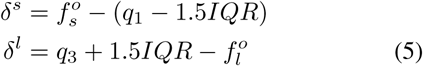
5. Append *I*^*o*^ with the index of the maximum, *i*^*o*^:

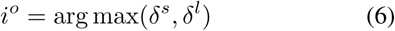
6. Go to step 2.

We used the IIQR algorithm on each beat for all features in the feature set. The final rejected beats were the union of those identified as outliers for each feature.

#### 3) Metrics

An expert inspected TCD recordings, annotated the beats, and labeled the beats. To assess the efficacy of the algorithm, we computed the clinically significant TCD parameters from the <u>average beat waveform</u> in each interval by considering (*i*) all the beats and (*ii*) only those identified using the new algorithm. Next, we compared these parameters with those identified from expert-accepted beats. Finally, we used the paired Wilcoxon signed-rank test to test the null hypothesis of whether the improvement from using the algorithm over using all beats was insignificant.

The following TCD parameters were computed from the average beat waveform, see Fig. 4 for illustration:

**Fig. 4.**
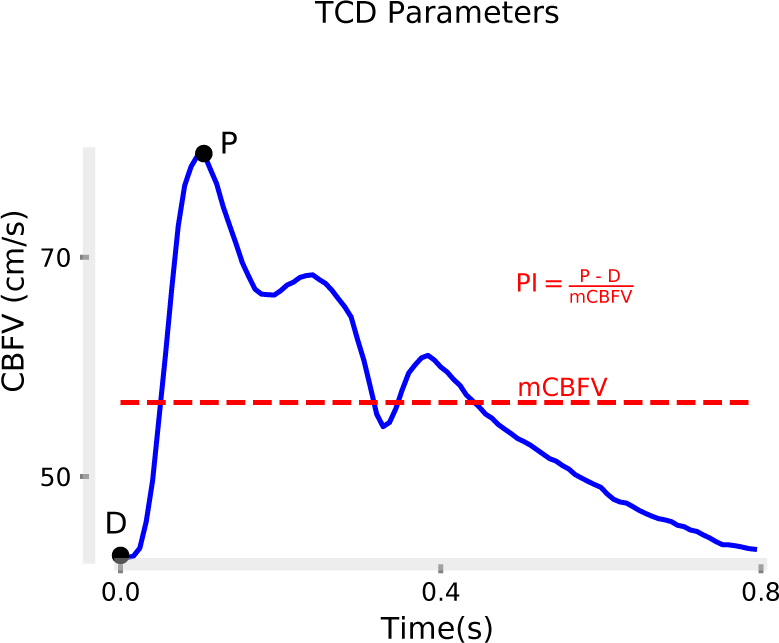
Typical average beat 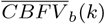 calculated in an interval. The figure is annotated to show the morphological points of the waveform that based on which the TCD parameters (e.g. mCBFV, PI) are defined.

- *Mean Cerebral Blood Flow Velocity* (mCBFV) is the mean of the CBFV waveform.

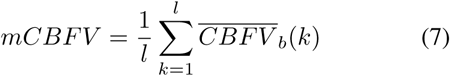
- *Pulsatility Index* (PI) is defined as:

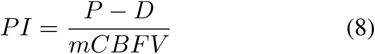

where *P* is the systolic peak and *D* is the diastolic valley. PI represents a measure of cerebrovascular resistance.

### C. Feature Selection

In order to identify the optimal combination of features proposed in II-B1, we systematically inspected each of the possible 31 combinations. To this end, we computed the improvement in percent error over using all beats *e*_*all*_ *-e*_*auto*_ in recovering the TCD parameters mCBFV and PI:

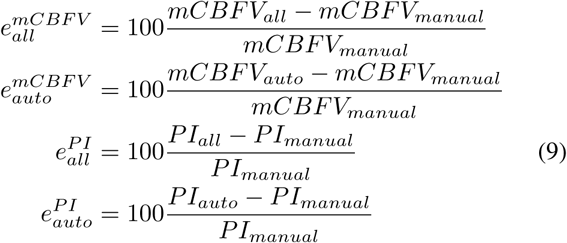

where (·)_*auto*_, (·)_*all*_, and (·)_*manual*_ refer to the TCD parameter identified from algorithm-accepted beats, all beats, and expert-accepted beats, respectively.

For the majority of recordings, improvement of using the algorithm-accepted beats over using all beats was positive.

For some recordings, however, the improvement was negative, which is because of frequent misclassified beats in a recording and/or misclassified beats that had a disproportionately large contribution to the TCD parameter mCBFV or PI. We quantified the performance of each feature combination using the sum of the positive improvements (*S*_*pos*_ green area under the curve in Fig. 5) and the sum of the negative improvements (*S*_*neg*_ absolute value of the red area under the curve in Fig. 5). Thus, the larger *S*_*pos*_ and smaller *S*_*neg*_, the more appropriate that feature combination. See Fig. 5 for illustration of the case when we use all five features. For the purpose of identifying the optimal combination, we consolidated the TCD parameters mCBFV and PI:

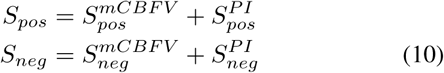

**Fig. 5.**
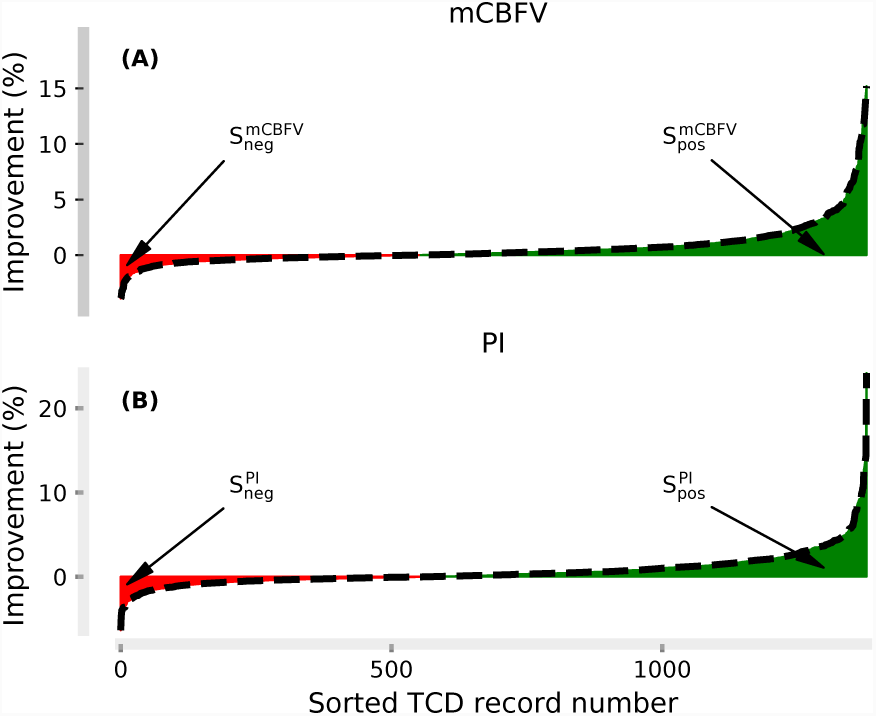
Sorted improvement gained by using the algorithm-accepted beats over using all beats in the computation of clinically important parameters: (A) *mean Cerebral Blood Flow Velocity* (mCBFV); (B) Pulsatility Index (PI). All five features ED-CC-BL-HFNP-DV were used for this figure. The area under the curve for positive improvement *S*_*pos*_ is shown in green and for negative improvement *S*_*neg*_ in red. The larger *S*_*pos*_ and smaller *S*_*neg*_, the more effective the algorithm in filtering TCD data to extract clinically important parameters.

To identify the variability of improvement for each feature combination, we performed a bootstrap analysis using 1000 iterations for each feature combination. At each iteration, we randomly selected the data with replacement and calculated the mean and variability of *S*_*pos*_ and *S*_*neg*_. We identified the optimal combination as the one with maximum *S*_*pos*_ *-S*_*neg*_.

## III. EXPERIMENTAL RESULTS

Fig. 6 illustrates the positive and negative improvements for the 31 possible feature combinations. There was a trade-off between *S*_*pos*_ and *S*_*neg*_, i.e. a feature combination with large *S*_*pos*_ (desired) typically resulted in larger *S*_*neg*_ (undesired). The best combinations, those which maximized *S*_*pos*_ – *S*_*neg*_, were CC and CC-DV. We could not reject the null hypothesis that the two best combinations had identical means (two-sided T-test, *p*-value > 0.05). The next best feature combinations were ED-CC and BL-CC, which were not statistically different from each other, followed by ED-CC-DV. The poorest feature combination was DV, followed by BL, HFN, BL-DV, BL-HFN. This suggests that for this study, cross-correlation may be the strongest feature in terms of identifying poor quality beats, as it was present in all five of the best performing combinations and was absent from the five worst combinations. However, there may be reason to believe that this result may not hold in general, which we explain further in Sec. IV-B. Perhaps more importantly though, is the fact that a large number of combinations containing different constituent features all perform significantly better compared with not performing any beat rejection. In fact, even the worst performing feature combination (DV) had a significantly larger positive improvement (*S*_*pos*_ = 850) over negative improvement (*S*_*neg*_ = 301). As seen in Fig. 6, the majority of other feature combinations comprise a significantly better performing cluster, which provides evidence for the robustness of the algorithm as a tool for generally differentiating between low and high quality segments of data, independent of the exact features selected.

**Fig. 6.**
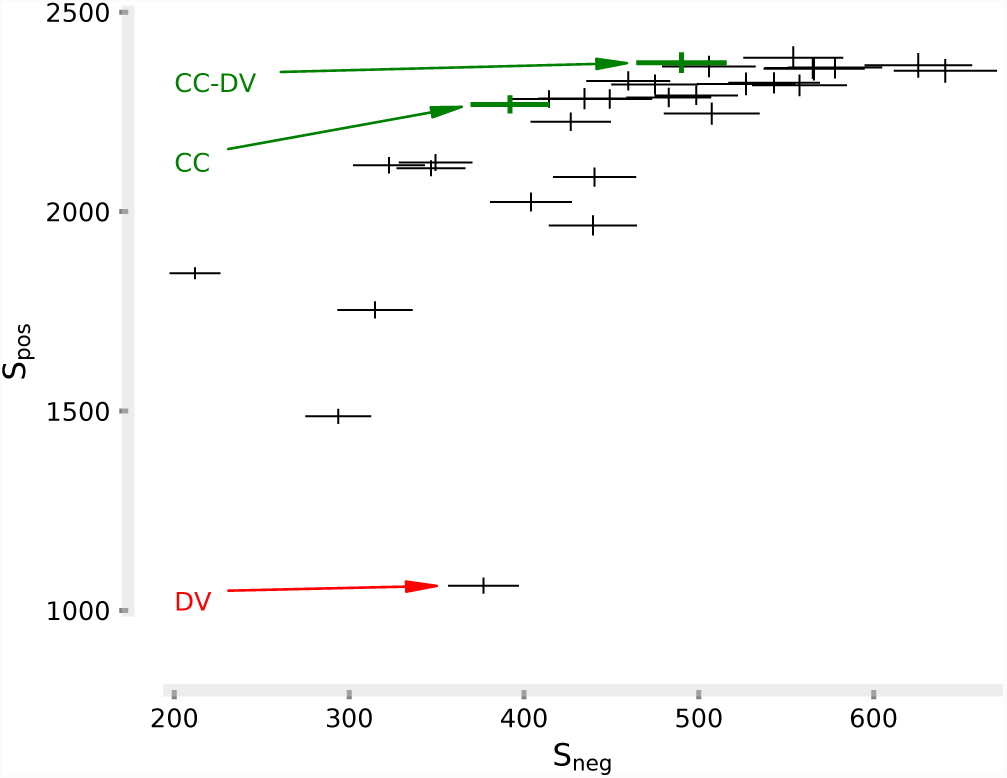
Cumulative positive and negative improvements *S*_*pos*_, *S*_*neg*_ for 31 possible feature combinations. Each combination is shown by a cross; the center of the cross represents the mean bracketed by vertical and horizontal bars representing standard deviations in *S*_*pos*_ and *S*_*neg*_ directions associated with the bootstrap study. The optimal combinations which maximized *S*_*pos*_ *-S*_*neg*_ were CC, and CC-DV.

Fig. 7 demonstrates typical performance of the beat rejection algorithm. It is evident that there is agreement between expert-accepted beats Fig. 7(A) and those accepted using our algorithm Fig. 7(B). In this example, however, there are three beats that are subject to disagreement; they are rejected by the expert but accepted by the algorithm. We elaborate on these disagreements in more details in Sec. IV-B. Fig. 8 visualizes the difference between average beat waveforms calculated from expert-accepted beats, algorithm-accepted beats, and all beats for the TCD recording shown in Fig. 7. The identified TCD parameters (mCBFV, PI) which are calculated from average beat waveforms were (38.8, 1.5), (38.2, 1.5), (33.7, 1.5) using expert-accepted, auto-accepted, and all beats respectively. Average beat waveform morphologies and the extracted TCD parameters demonstrate great consistency between the algorithm and expert. Evidently, the beats that were the subject of disagreement had little effect on the final average beat waveform morphology and thus on the extracted TCD parameters. However, the difference between all beats and expert-accepted beat waveforms was quite large, highlighting the importance of filtering out poor quality beats in order to acquire more accurate TCD parameters.

**Fig. 7.**
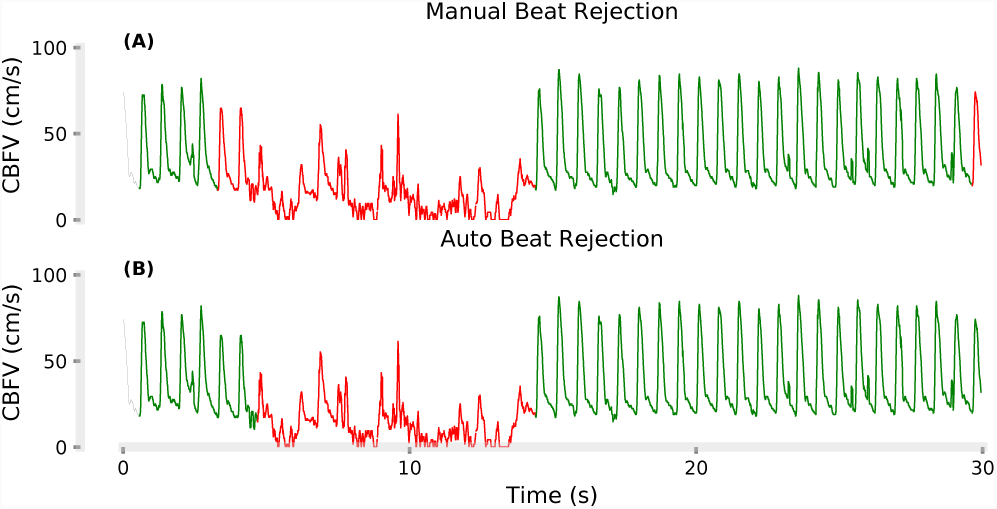
Typical TCD recording annotated by an expert (A) and by the beat rejection algorithm (B). Red represents the rejected beats and green the accepted beats.

**Fig. 8.**
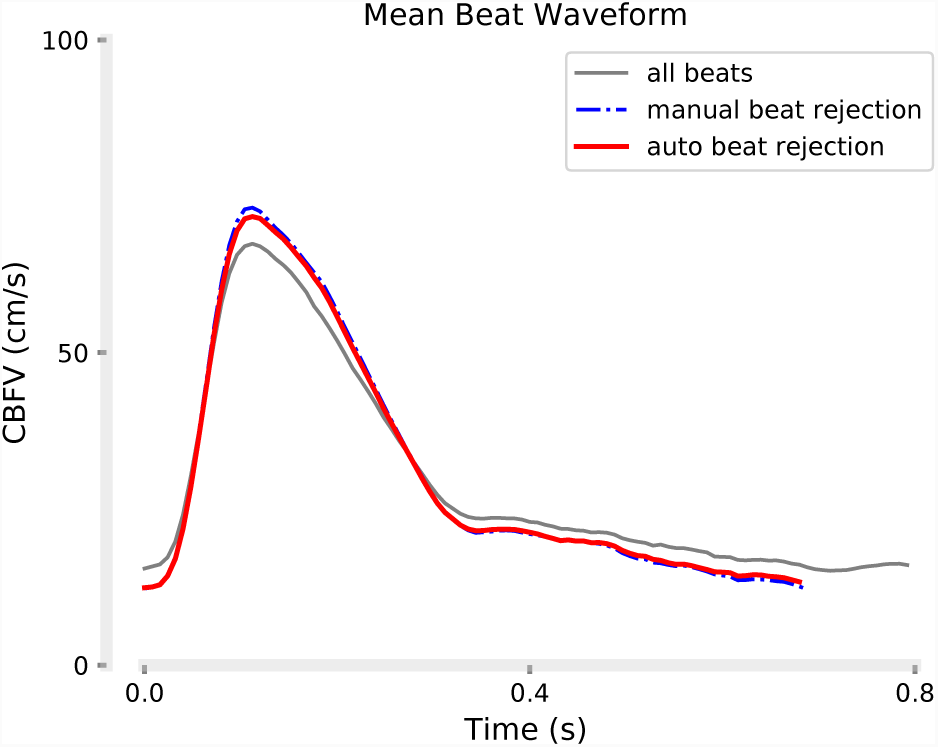
Comparison of the average beat waveform from all beats accepted, expert accepted beats, and algorithm accepted beats for the TCD recording shown in Fig. 7. There is great consistency between the expert and the algorithm. The identified TCD parameters (mCBFV, PI) were (38.8,1.5), (38.2,1.5), (33.7,1.5) using expert-accepted, auto-accepted, and all beats respectively

Fig. 9 and 10 show the group results. To compare error distributions, Fig. 9 depicts the empirical *Cumulative Distribution Functions* (CDF) for percent errors using the algorithm-accepted beats *e*_*auto*_ and all beats *e*_*all*_. The CDF plots show that the error range is smaller and the CDF approaches unity much faster for auto-accepted beats. Consequently, smaller errors are more likely when using the algorithm. Fig. 10 demonstrates that the higher the error from using all beats, the more likely the algorithm is to achieve a higher improvement, and that there is a statistically significant and strong positive correlation between the two. Moreover, the signed-rank test between *e*_*auto*_ and *e*_*all*_ rejected the null hypothesis that the population means were the same (*p*-values < 10^-4^).

**Fig. 9.**
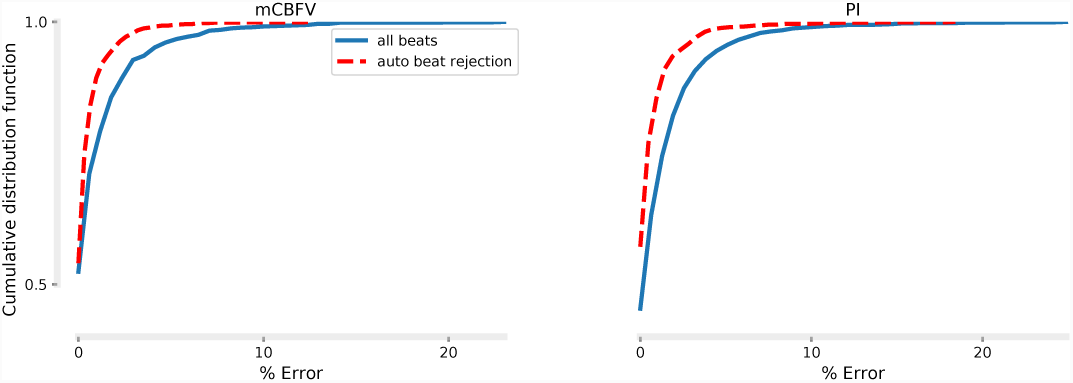
Cumulative distribution functions of percent errors in estimating the TCD parameters: (A) mCBFV and (B) PI using auto-accepted and all beats. The error range is smaller when using the auto-accepted beats and the CDF reaches unity faster which demonstrates the utility of using the algorithm.

**Fig. 10.**
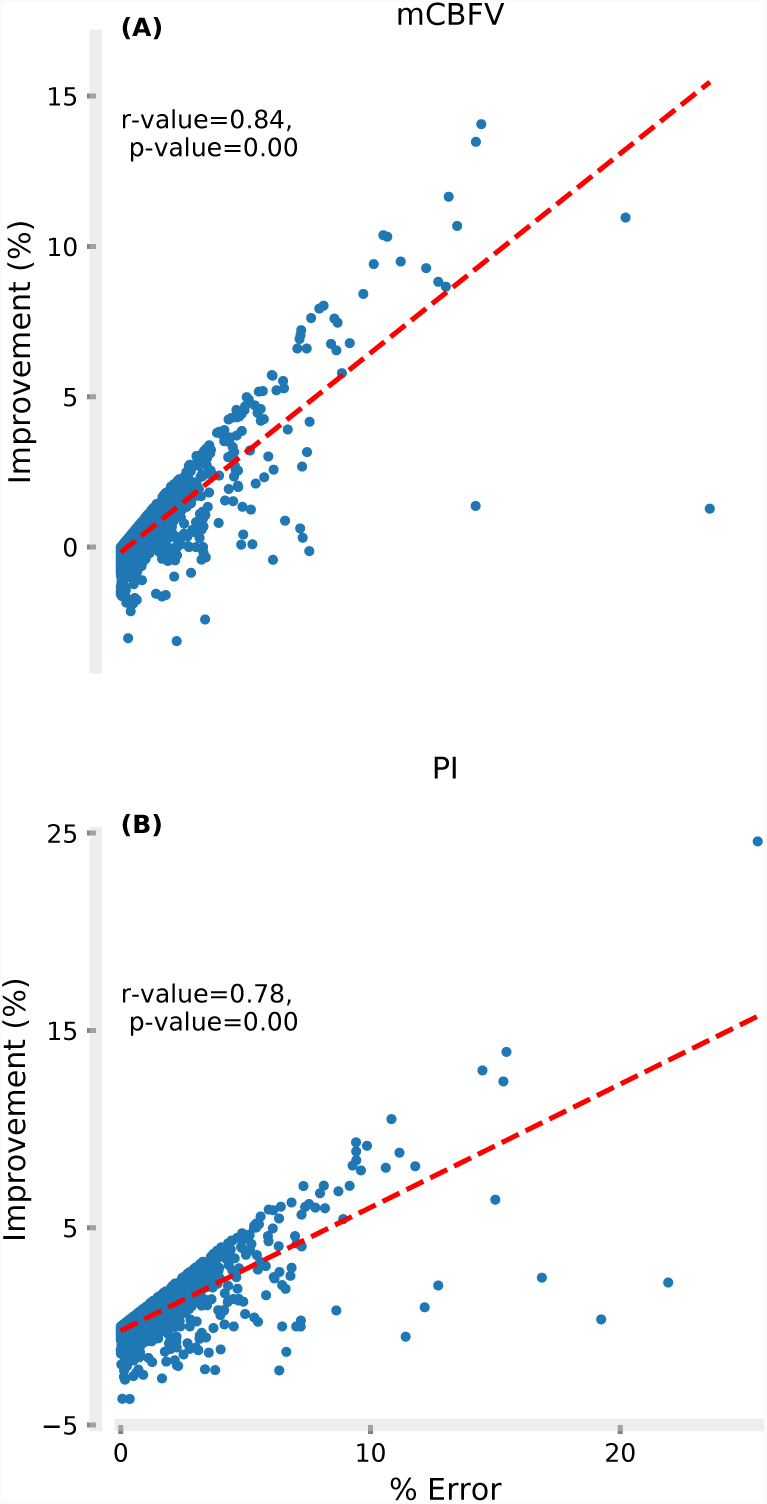
Scatter plots of improvement in estimating the TCD parameter using auto-accept beats as a function of error using all beats show statistically significant, strong positive correlations.

## IV. DISCUSSION

### A. Conclusion

In this paper, we proposed objective features to assess beat quality in TCD recording from cerebral arteries. We described the *Iterative InterQuartile Range* (IIQR) algorithm, an outlier rejection method that used these features to label beats that were different from others in a TCD recording. We assessed the performance of this tool on a dataset consisting of over 16 hours of TCD data with more than 76,000 beats recorded from healthy and pathological subjects. We identified that the best feature for identifying poor-quality beats was cross-correlation for our data. We demonstrated that the IIQR algorithm successfully identified low-quality beats and the clinically important TCD parameters from the algorithm-accepted beats were consistent with those computed from beats accepted manually by an expert.

### B. Data

In order to properly interpret the results of this study, a number of facets pertaining to the data require discussion, specifically with regard to the expert annotations. A major part of the motivation for this work is the inherently subjective and unrepeatable nature of manually evaluating data quality. This limitation persists even for the most experienced TCD experts, and it is important to keep this in mind when drawing comparisons to the expert-accepted beats. Furthermore, while the difference between highly poor and good beat is generally self-evident, no objective classification criteria exists. An unavoidable consequence of this is low inter-and intra-rater reliability for intermediate range of signals. As a result, no true gold standard can exist for this type of work, which can complicate the task of evaluating the IIQR algorithm’s performance. Consequently, disagreement between the algorithm and expert does not imply inaccuracy and isolated disagreements can be simply the side affect of rater variability and lack of objective classification criteria.

Nevertheless, some method of evaluation is required, and some metric of performance, however imperfect, is needed. It is clear from the examples shown here, and based on inspection by other independent experts that the set of manually accepted beats used in this study represents a significant improvement in signal quality over simply using the set of all detected beats. While exact agreement is not necessarily desirable or even achievable, the error function provides an objective way of quantifying the discrepancies to compensate for the lack of a gold-standard. Finally, to facilitate the manual inspection process, a software tool was developed to display the data along with the detected beats, which would allow the user to add, delete, or shift beat start and stop indices. In addition, this tool included features to aid the user in identifying potentially bad beats by flagging beats that were abnormal according to various metrics (which included cross correlation) relative to the other beats in the scan. While the user wasultimately responsible for manually labeling the data, the features used to automatically highlight probable beat outliers could have served as a bias during the inspection process, and results should be viewed through this lens. This fact is especially important for discussions involving the relative importance or performance of individual features. Nevertheless, we feel that one of the more important takeaways from this work is that any combination of the features discussed resulted in significantly improved performance, a result which appears to be robust in spite of the limitations or biases present in this study.

### C. Features & Algorithm

To ensure fidelity of reported clinical parameters from TCD measurement, an expert typically inspects the data and labels/removes the low-quality portion of the data, a process which is fraught with problems, as previously detailed. Quantitative measures of TCD signal quality are of paramount importance for the accurate extraction of clinical parameters. However, the TCD beat waveform is complex and can change based on subject, age, race, health, vessel, and sonographer skill, among other things. Therefore, absolute feature criteria for beat classification can be inappropriate as they can change from recording to recording independent of data quality. Consequently, we chose to classify beats by comparing their features relative to others in a recording and developed the IIQR algorithm to identify those with substantially different features.

In this work, we have presented quality features for each beat to be compared against those from other beats in a recording for decision making. Some of these features are based on the difference between a beat waveform and the average beat waveform in the recording including Euclidean Distance and Cross-Correlation. These two metrics are meant to identify beats whose waveform morphology is significantly different. The rest of the features were beat length, high-frequency noise power, and diastolic variance. Deviations in beat length can be attributed to both the inherent healthy and pathologic variability of the heart. More importantly, beat length deviations can also be due to errors in identifying the beat onset and end point. It is also common to fail to detect any beats in portions of very low quality signal, resulting in abnormal lengths. Thus, it is in our interest to label beats that are either outliers in terms of beat length, i.e. statistically longer or shorter than their adjacent beats. Inclusion of these beats may compromise the quality of the average beat waveform. High frequency noise power ratio can be correlated with signal-to-noise ratio and is a measure of undesired high-frequency noise in the acquired signal since the CBFV should have power primarily in low frequencies only (*f* < 15*Hz*). Diastolic variance is a measure of variability in the diastolic portion of the beat which is expected to have low variability but occasionally has high variability due to noise. These features were chosen based on consulting with experts in TCD signal recording and interpretation.

With the five proposed features, we had 31 different combinations of features to choose from. In order to find the optimal feature combination, we systematically assessed each feature, inspected the results and compared the clinically important parameters against those obtained from expert-accepted beats. We identified a trade-off in selecting features for beat rejection: the higher the positive improvement for a combination, the higher the negative improvement which is visualized in Fig. 6. Upon inspection of the classifier performance, we identified that the combinations with many features typically resulted in larger positive improvements when quality was poor and many beats needed to be rejected. However, these combinations also rejected beats in recordings where beat quality was not poor and resulted in negative improvements. Therefore, we selected the combination that maximized the “score function” defined as the negative improvement subtracted from positive improvement. We identified that cross-correlation was the best feature to make beat rejection decisions. It is important to note that this choice was optimal only for our labeled data. Theoretically, this choice could be different depending on the experimental conditions, for example with a different ultrasound probe, data acquisition system, sonographer, etc. Nevertheless, as a general rule of thumb, we believe that cross-correlation is a good first feature to inspect. In addition, if there are strong a priori reasons to choose certain features based on knowledge of the data or particular study, then the appropriate features can be selected.

We have developed the *Iterative InterQuartile Range* (IIQR) method to detect outlier beats. We chose IQR because it is non-parametric, robust to outliers, requires no *a priori* information about the data, and is not guaranteed to always label a portion of the data as outliers. Outlier detection algorithms typically work by comparing a data point to the statistics gathered from all the available data points to make a decision about whether the data point is an outlier. Importantly, the outliers themselves are used to compute the statistic, which under the right circumstances, can bias the resulting statistics enough to prevent the detection of all outliers. To get around this problem, we chose to develop an iterative algorithm to recompute the statistics each time an outlier is removed. At each iteration, we remove only the most prominent outlier and recompute the statistics, iterating until no further outlier can be detected. While this approach is more accurate in the computation of the statistics due to minimizing the contribution of outliers in the statistics, it can be computationally more expensive due to the number of the iterations.

The IQR method is a two-sided outlier rejection technique and identifies both too low and too high outliers. However, for some features, it is more appropriate to have a single-sided rejection rule. Thus, we also developed and used the single-sided rejection versions of the IQR method as needed. For high frequency noise, Euclidean distance, and diastolic variance, we only rejected too high outliers as low values for these features are desired by definition. For cross-correlation, we only rejected low outliers as high correlation is desired by definition. For beat length, however, we used the two-sided version as both low and high beat lengths can be inappropriate.

The final rejected beats were the union of beats identified by each of the features since each feature inspected a different physiological aspect of the beats. While there was overlap between the features, this is a simple and the least conservative approach. In future, it is of interest to study more complex feature voting algorithms.

Since the proposed beat rejection mechanism is based on outlier detection, it can possibly break if the overall data quality is low. For example, the algorithm will likely pass all the poor beats if they are the majority. Future work is needed to remove beats altogether in a second stage based on absolute beat features to ensure that a minimum TCD signal quality is retained.

### D. Other Algorithms

There are a few other relevant algorithms that can be used to assess and improve the quality of the TCD signal. Gunn’s recent technique requires an auxiliary signal [19]. They have developed a system identification algorithm and trained a model between *Arterial Blood Pressure* (ABP) signal and CBFV which detected and corrected for artifacts in CBFV. It assumed that artifacts are sparse events that increase the complexity of the dynamics of the system between ABP and CBFV. This approach requires measurements of additional signals i.e. ABP. Moreover, the identified model will presumably need to be tuned for each TCD recording to account for intersubject and intrasubject variabilities and differences in the waveform between health and disease.

Similar to TCD recordings, Intracranial Pressure (ICP) is a triphasic pulsatile waveform that can be contaminated with noise and artifacts. A few methods have been developed to improve the ICP/CBFV data quality as well as the extraction of additional features [20], [21]. As one example, the MOrphological Clustering and Analysis of Intracranial Pulses (MOCAIP) algorithm utilizes a five-step process, including a hierarchical clustering method to construct a representative non-artifactual beat known as a dominant pulse [20], [22], [23], [24], [11]. Following the clustering of the pulsatile beats, the dominant pulse is compared to a pulse library to determine if it is a spurious pulse and sub-peak landmarks are identified for post-hoc feature extraction. Although this work was originally used for ICP pulses, it was later adapted for the analysis of CBFV waveforms [11]. In comparison to the IIQR algorithm developed in this work, there are a few advantages and disadvantages. First, the identified clusters in the MOCAIP algorithm could be meaningful, e.g. cluster with pathologic pulse waveform or high-noise cluster, etc. Second, the use of the validated pulse library in MOCAIP provides insight into the quality of the signal collected, e.g. if there is no signal, the MOCAIP algorithm will not return valid results. However, it is this pulse library and post hoc analysis that does not allow MOCAIP to be used in a near real-time use, such as within the clinical environment for fast diagnostic purposes where the IIQR could be. Furthermore, the IIQR algorithm is more general in the sense that we can label and reject beats based on a select feature (five proposed in this paper) whereas MOCAIP is based on Euclidean distance between the beats.

### E. Clinical Implication

This paper presents a tool that can be used as a prefiltering stage to improve data quality for assessment of many cerebrovascular pathologies including *Acute Ischemic Stroke* (AIS), [25], [3], [26]. The limited number of available effective treatments and interventions are highly time-sensitive. Therefore, to avoid unnecessary delays, pre-hospital diagnostic tools are critical for the fast triage and transfer of patients to specialized and appropriate treatment centers. Because TCD measures blood flow through the cerebral vasculature directly in a fast, portable, and noninvasive way, it is a strong candidate technology for pre-hospital diagnosis and assessment [27]. Furthermore, analysis of subtle, clinically significant changes in CBFV waveform morphology has shown promise for potentially discovering new biomarkers and developing machine learning algorithms to aid in performing objective diagnostic assessments for diseases which affect the cerebrovasculature [13], [14], [28].

Nevertheless, difficulties associated with acquiring high quality TCD data have hindered its adoption for this purpose. Due to the ability of the algorithm presented here to reliably assess the quality of local segments of data based on objective criteria, we have taken yet another step forward toward alleviating the obstacles preventing the adoption of TCD as a pre-hospital diagnostic tool. In being able to quickly and accurately extract the high quality portions of signal from a TCD recording, this algorithm may be able to aid in diagnostic assessments for conditions such as AIS as well as many other cerebrovascular conditions, which often rely heavily on obtaining an accurate, clean picture of the beat waveform present in the signal. Significantly, the algorithm was able to closely replicate the performance of a TCD expert in terms of measured clinical parameters, yet it has none of the disadvantages normally associated with manual inspection.

## VI. ACKNOWLEDGMENT

The authors thank Dr. James K. Fleming, Ms. Brenda Knowles, Mr. Leo Martinez, Mr. Ben McClellan, Ms. Jennifer Nichols, Ms. Jennifer Patterson, and Dr. Ruchir Shah for their help during data collection and subject recruitment.

